# X-DPI: A structure-aware multi-modal deep learning model for drug-protein interactions prediction

**DOI:** 10.1101/2021.06.17.448780

**Authors:** Penglei Wang, Shuangjia Zheng, Yize Jiang, Chengtao Li, Junhong Liu, Chang Wen, Atanas Patronov, Dahong Qian, Hongming Chen, Yuedong Yang

**Affiliations:** Institute of Medical Robotics, Shanghai Jiao Tong University, Shanghai, China; Guangzhou Regenerative Medicine and Health Guangdong Laboratory, Guang-zhou 510530, China; School of Data and Computer Science, Sun Yat-sen University, Guangzhou, China

**Keywords:** drug-protein interactions, virtual screening, graph neural network model, protein graph, deep learning

## Abstract

**Motivation:** Identifying the drug-protein interactions (DPIs) is crucial in drug discovery, and a number of machine learning methods have been developed to predict DPIs. Existing methods usually use unrealistic datasets with hidden bias, which will limit the accuracy of virtual screening methods. Meanwhile, most DPIs prediction methods pay more attention to molecular representation but lack effective research on protein representation and high-level associations between different instances. To this end, we presented here a novel structure-aware multi-modal DPIs prediction model, X-DPI, performing on a curated industry-scale benchmark dataset.

**Results:** We built a high-quality benchmark dataset named GalaxyDB for DPIs prediction. This industry-scale dataset along with an unbiased training procedure resulted in a more robust benchmark study. For informative protein representation, we constructed a structure-aware graph neural network method from the protein sequence by combining predicted contact maps and graph neural networks. Through further integration of structure-based representation and high-level pre-trained embeddings for molecules and proteins, our model captured more effectively the feature representation of the interactions between them. As a result, X-DPI outperformed state-of-the-art DPIs prediction methods and obtained 5.30% Mean Square Error (MSE) improved in the DAVIS dataset and 8.89% area under the curve (AUC) improved in GalaxyDB dataset. Moreover, our model is an interpretable model with the transformer-based interaction mechanism, which can accurately reveal the binding sites between molecule and protein.

## 1 Introduction

The identification of drug-protein interactions (DPIs) lies at the core of in silico drug development. Though experimental assays remain to be the golden standard for determining the binding affinities and modes, experimental characterization of every possible drug-protein pair is daunting as there are over 166 billion drug-like compounds [1] and over 5000 potential protein targets [2]. Alternatively, hit compounds could be identified for given protein targets effectively and inexpensively through computational approaches.

Many computational methods have been developed, and these methods could be generally split into two categories: physic-based and machine-learning methods. Physic-based methods like molecular docking apply physics-inspired force fields to simulate the binding of a protein and a molecule at the atomic level and to estimate the binding free energy between them [3]. However, the performance of these methods is often unsatisfactory due to the difficulty in assessing the solvent contributions and conformational entropy. In addition, these physical methods are sensitive to structural fluctuations, which prevents them from dealing well with the flexibility of proteins [4].

On the other hand, machine learning-based methods recently met rapid progress with the recent increase in the available protein-ligand binding data and a decrease in computational costs [5-7]. The general idea for this method is to integrate structural information from ligands, proteins, and their interactions into a unified framework. In this case, molecules can be characterized by molecular fingerprints, structural descriptors, or topographies, while proteins can be described by sequences or tertiary structures. Their representations are then extracted by the designed neural network to obtain abstract information and are eventually used to predict whether and how they will bind to each other [5, 8-12].

Despite a lot of previous efforts, there are a few general caveats among these proposed models:

### Using unrealistic datasets with hidden bias

Although a large amount of experimentally reported structure-activity relationship (SAR) data is available, data collation and cleaning are quite tedious and laborious processes. Current deep learning works either used very limited datasets, such as the Human dataset and C.elegans dataset [7] which include positive DPI pairs from DrugBank 4.1[13] and Matador [14], and highly credible negative samples from a systematic screening framework [15], or use arbitrary benchmark with expert-defined *decoys* (i.e., negative samples were generated by fixed rules), including DUD-E, MUV [16] and so on. These datasets are unfortunately biased by both obvious and hidden chemical biases, therefore overestimating the true accuracy of virtual screening methods. For example, DUD-E was collected with the intention to train structure-based virtual screening, and thus the ligand-based split is extremely native for this benchmark. These datasets can be easily separated by ligand information and cannot guarantee that models learn protein information or interaction features. Instead, the key usage of DPI is to identify hit compounds for unseen and non-homolog with few known actives. In addition, the datasets from high throughput screening (HTS) tend to be much larger and noisier, and most of the currently delicate and fragile models might not be robust enough to deal with these real-world data.

### Suboptimal representation of protein

Although the key of DPI is the generalization of unseen and non-homolog with few actives, current works [5, 10-12] normally use one-hot encoding vectors to represent residues. This conventional approach neither is able to embed the contextual dependencies between residues nor to make use of protein topological information. In fact, the protein topological information is crucial information for determining the binding affinity between protein and drug in practice [17] and several methods [4, 18] have shown that the protein structural information like 2D distance map is an effective feature for DPI prediction. The simple sequence representation of protein cannot even capture the connection between different proteins, let alone the binding mode between protein and drug. Although the direct input of 3D structure has been introduced in recent studies [6, 19, 20], they cannot address the issue of 3D transform invariance properly. Moreover, the lack of protein structure hampers the development of real structure-based protein representation models in the DPI task.

### Lack of the high-level associations of instances

Existing deep learning models mainly focused on the information of the input drug-protein pairs, ignoring the high-level information from protein-protein associations (PPAs) and drug-drug associations (DDAs). The significance of PPAs and DDAs derives from a well-established hypothesis that proteins typically bind with similar drugs [21], which is key to generalizing DPI predictions. Earlier studies generally considered association by using molecular fingerprinting [22] techniques or BLAST [23] to calculate the similarity of co-evolutionary information. However, these approaches were limited in dealing with homologous proteins and had difficulties in dealing with unseen proteins and drugs with novel scaffolds.

### Present Work

To address these challenges, we propose a novel structure-aware multi-modal method (coined as X-DPI) for in silico DPI prediction. X-DPI is enabled by the following contributions:

1. We curated a large-scale benchmark GalaxyDB specifically designed for machine learning-based virtual screening. GalaxyDB was derived from ExcapeDB and consists of 372 common targets with 381,021 confirmed active and 1,634,038 confirmed inactive compounds. The large data size and an unbiased training procedure provide advantages for model building than using a small toy dataset.
2. For informative protein representation, we constructed a structure-aware graph neural network method from the protein sequence by combining predicted contact maps and graph neural networks.
3. We introduced self-supervised pre-trained embedding of drugs and proteins, respectively, in order to strengthen the protein/drug association signals. Our model leverages this high-level information in a unified framework and generates interpretable results with a transformer-based interaction mechanism.

We provided a comprehensive performance comparison among several state-of-the-art (SOTA) methods. Our results demonstrated that X-DPI has superior performance over some SOTA models by up to 8.89% improvement. In addition, we also made a prospective prediction on Davis dataset [24] and the external dataset extracted from the AstraZeneca screening database.

## 2 Methods

### 2.1 Dataset construction

We constructed experiments using the following two benchmark datasets for model building and evaluation. In addition, an external test dataset from AstraZeneca was utilized to verify the generalization ability of our model on the industrial dataset.

a. **Davis dataset** consists of binding affinity information with *K*_*D*_(kinase dissociation constant) values among 72 drugs and 442 targets. In our experiments, we use SMILES representation of 68 drugs and sequence representation of 442 target proteins from the DeepDTA [11] training/test dataset. For the Davis dataset, we view the DPI prediction task as a regression task that predicts the *K*_*D*_values for each DPI pair. This small dataset is used to initially verify that our model can effectively deal with the DPI prediction problem. When we use the Davis dataset for prospective validation, we assign the data points in the Davis dataset to two class according to the criterion of *K*_*D*_≥6 and view the DPI prediction task as a classification task.
b. **GalaxyDB**. We curated a large-scale DPI benchmark, GalaxyDB, based on the ExCAPE-ML [25], a collection of protein-ligand entries complied from ExCAPE-DB. ExCAPE-ML is composed of 955,386 compounds, covering 526 distinct target proteins for a total of 49,316,517 structure-activity relationships (SAR) data points. For classification tasks, the data points were assigned to two classes (i.e., inactive, active) according to their log-transformed activity values (pXC_50_ values). A compound-target record was defined to be activated if it fulfilled the criterion of pXC_50_≥6 (activity≤1µM). We analyzed the distribution of the original ExCAPE-ML dataset for the regression task and observed that there are a large number of (more than 45 million) data points with a pXC_50_ value of 3.101 in the dataset, which are expert-defined negative samples with low confidence. For curating a high-quality dataset, we excluded the data pXC_50_ value of 3.101 on the basis of the ExCAPE-ML dataset to form a relatively balanced and high confident benchmark set. Subsequently, the dataset was trimmed down by removing the target proteins with a sequence length longer than 750 in order to reduce the computational cost during the calculation of the contact map and co-evolution features for proteins and the processing of protein features. Finally, we have obtained a high-quality benchmark dataset, GalaxyDB with an appropriate quantity, which is composed of 632,459 compounds covering 372 distinct target proteins and in a total of 2,015,059 DPI data points. We selected this benchmark dataset for the training and evaluation of our proposed model.
c. **External test dataset**. To test whether the model is capable of performing real-world virtual screening tasks, we have done prospective prediction with AstraZeneca in-house SAR data. In particular, for targets seen in our train set, we selected the top 30 targets according to the performance of our model in the GalaxyDB dataset and required that each target need to have at least more than 100 data points. Additionally, we also selected 10 targets that are not included in our training set. In total, we constructed an external test set which is composed of 208,958 data points, including 172,768 data points for 30 targets seen in the training set and 36,190 data points for 10 unseen targets.

### 2.2 Representation of Protein and Molecule

The representations of protein and molecule lie at the core of the DPI task. In this section, we described the initial feature representations of target proteins, followed by the feature representations of molecules.

#### Protein representation

The protein representation was done from the perspectives of structure and sequence features, respectively. For the structural feature, we considered using a graph to represent the 2D structure of proteins, which has been proven effective for predicting protein solubility in our previous study [26]. In the protein graph model, residues were regarded as nodes and the contact map predicted from the sequence was used as the adjacency matrix. Here node features were represented by the Hidden Markov Matrix (HMM), position-specific scoring matrix (PSSM) and structural features predicted from *SPIDER3* [27]to represent the node features and used the predicted contact map to represent the adjacency matrix in protein graph. The PSSM and HMM features are evolutionary information that contains the motifs related to protein properties in protein sequences [28]. And the PSSM profile was generated by *PSI-BLAST v2*.*7*.*1* [23] with the *UniRef90* sequence database after 3 iterations, the HMM profile was generated by *HHBLITS v3*.*0*.*3* in aligning the *Uni-Clust30* profile HMM database [29] with default parameters. The structural features include 14 features to reflect the secondary structure of proteins predicted by *SPIDER3*. The list of protein node features can be found in the supplementary material. For the contact map of proteins, we made predictions of the protein contact map by SPOT-Contact [30], which takes the protein sequence-based and evolutionary coupling-based information as input to predict the contact probability of all residue pairs in one protein. Finally, we obtained a protein graph as *G*_*a*_ = (*V*_*a*_, *A*_*a*_), where *V*_*a*_ ∈ *R*^*n*×*f*^ is the set of *n* amino acid nodes, each node represented by *f*-dimension features vector composed of HMM, PSSM and structural features, *A*_*a*_ ∈ *R*^*n*×*n*^ is the adjacency matrix (contact map) for the protein graph.

For the protein sequence feature, we also considered using the high-level representation learned from a large collection of unlabeled protein sequences provide by TAPE [31]. TAPE is a language model for protein representation and it encodes each amino acid into an embedding vector. For each embedding vector, it is contextual and includes the sequence information from the input protein sequence, so we embedded the protein sequence to tape embedding with the pre-trained BERT [32] model in TAPE.

#### Drug Molecular representation

We represented the drug molecule as a graph to get more accurate structure information for the molecule. In this sense, a molecular graph can be formulated as *G*_*c*_ = (*V*_*c*_, *A*_*c*_), where *V*_*c*_ ∈ *R*^*n*×*f*^ is the set of *n* atom nodes with each node represented by *f*-dimension features vector composed of atomic properties. Here we used *f*-dimension atomic features that are detailed in the supplementary material. *A*_*c*_ ∈ *R*^*n*×*n*^ is the set of edges represented by the adjacency matrix for the molecular graph. The existence of edges in the adjacency matrix depends on whether the corresponding atoms in the molecule directly have a covalent chemical bond. Besides, we also used mol2vec [33] features at graph level as a high-level representation for a molecule to capture the DDAs.

We believe that the additional high-level representation from pre-trained embedding for proteins and molecules could provide implicit information to make the model distinguish different proteins and molecules. The high-level representation provides the global similarity information for DPI models, which describes the protein-protein association and drug-drug associations (PPAs and DDAs). The DPI models could leverage the global similarity to measure the associations between the seen and unseen proteins and molecules and make full use of the features of existing data to improve the performance of DPI prediction.

### 2.3 Model architecture of X-DPI

The overview framework of our proposed X-DPI network is shown in Figure 1. The input information includes multi-level representation for proteins and ligands. As shown in Figure 1, our model consists of three main modules: a graph representation network for proteins (Protein GNN Encoder), a graph representation network for molecules (Molecular GNN Encoder), and an interactive network with a transformer decoder for message interaction (Interaction Decoder).

**Figure 1.**
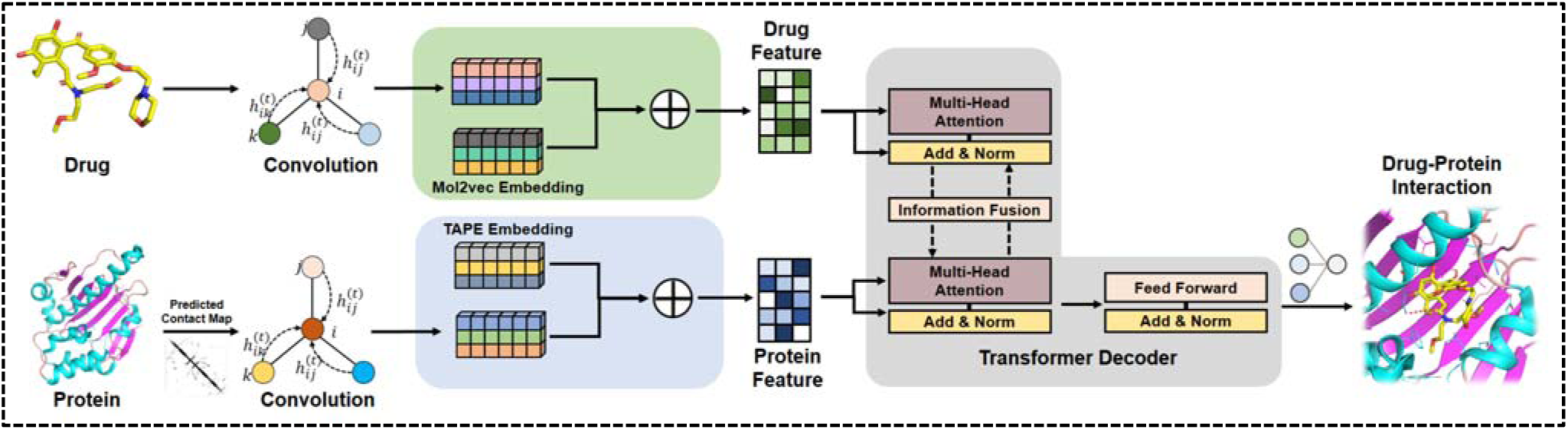
The architecture of X-DPI. It first processes the molecule and protein features parallel, then fuses the embedding of molecule and protein by Transformer Decoder for the DPI prediction.

#### Protein GNN Encoder

In our model, graph representation of proteins and the pre-trained feature encode by TAPE were input to the Protein GCN Encoder to learn structure and sequence representation of proteins at the same time. The Protein GCN Encoder includes two aspects: the first is the GCN encoder which encodes the structure information of protein graphs. The second is an information fusion unit to fuse the embedding information from the GCN encoder and the high-level representation from the pre-trained model. The protein graph was represented as a combination for the contact map and node features as input for the GCN encoder, then the GCN encoder learns node-level outputs for the protein graph. The GCN can be used to effectively process the graph structure data. The propagation rule can be represented in the normalized form as Equation 1:

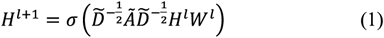

where Ã = *A* + *I*_*N*_ is the adjacency matrix of the graph with added self-connection. 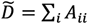 which is the diagonal node degree matrix. *W*^*l*^ and *H*^*l*^ are the learnable parameters in GCN and the output of *l*-th layer respectively. σ(·) is an activation function such as ReLU. For protein graph, *H*^0^ = *V*_*a*_, *A* = *A*_*a*_. To further extract the high-level features for protein, we also used CNN with Conv1D and gated linear unit (GLU) to fuse the different node embedding. In addition to the structure information from the protein graph, we also used dense layers to encode the extra protein sequence embedding information generated from the TAPE model. Finally, the protein graph and sequence information were concatenated to form the protein feature for the following Inactive Decoder module.

#### Molecular GNN Encoder

Similarly, the GCN was also used to encode the molecular graphs. In particular, we used the same GCN architecture as in the Protein GNN Encoder to learn the node-level features for molecules and obtain the molecule structure embedding. On the other hand, the dense layers were used to encode the high-level representation information from the mol2vec embedding, then the structure and mol2vec embedding information were concatenated to form the molecule feature for the following Interaction Decoder module.

#### Interaction Decoder

This module we used was inspired by the TransformerCPI [8], which provides a method to fuse the embedding features of molecule and protein. A transformer decoder was leveraged in our Interaction Decoder module to combine the information of proteins and molecules. The Interaction Decoder here served as a fusion unit to capture features useful for the interaction between molecule and protein. The decoder mainly consists of a multi-head self-attention layer and feedforward layer. The multi-head self-attention layer employed the multiple self-attention mechanisms to extract interaction information and it can be represented as:

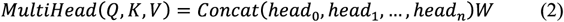

where the *Q, K* and *V* are the queries, keys and values in the Transformer. *head*_*i*_ is the output of *i*-th self-attention layer, 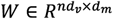 is a learnable parameter for fusing attention information from different heads. *n* and *d*_*m*_ are the number of heads and the dimension of the hidden state respectively. The self-attention in each head calculates the attentions by:

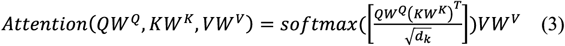

where the projection matrices 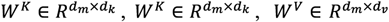 are learnable parameters. Compared with previous methods of directly concatenating the protein and molecule embedding information, we believe that this architecture can more effectively capture the interaction between protein and molecule embedding. Finally, we obtained the interaction features between protein and molecule and we could calculate the molecular interaction with a protein as follows:

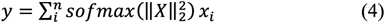

where *X* is the output matrix of transformer decoder and composed of a set of interaction vectors *x*_1_, *x*_2_, …, *x*_*n*_. The 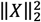 represent the *l*_2_ norm for *x*_*i*_ in interaction matrix *X. n* is the number of a set of interaction vectors from the Transformer Decoder. Finally, the interaction feature *y* was fed into a fully connected layer and a sigmoid function and obtained the predicted interaction probability 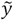 between protein and molecule. The model would be trained by maximizing the likelihood of regressing the training data, which means minimizing the binary cross-entropy loss as follows:

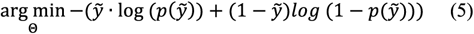

where Θ are the learnable parameters of the model.

### 2.4 Model training and evaluation

Our model took protein graphs, protein evolutionary and predicted structural features, molecule graphs, molecular substructure features as input, where we converted the SMILES representation for the molecule to graph representation through RDKit [34]. The model was implemented in Pytorch and trained on RTX 2080Ti. And the training details for our model as follows: The hidden state size *d*_*m*_ were set to 64 and 256 for molecule embedding and protein embedding, respectively. The number of graph convolution iteration was set to 3, and the kernel size in CNN for protein embedding was 7. For Transformer Decoder, the number of decoder layers was set to 3, and the number of heads in the multi-head layer was set to 8. Apart from all the hyper-parameters mentioned above, the maximum number of epochs during the training process in our model was set to 50, as the performance no longer improves after 5 epochs in the validation dataset for the classification task and the batch size equals 32 in every epoch. For the regression task, we trained the model with 1000 epochs and select the best model by RMSE in the validation dataset. Dropout was applied in CNN and the Transformer Decoder layer and the dropout rate is set to 0.2. For the optimizer in our model, we used the LookAhead [35] optimizer combined with RAdam [36] optimizer, in which the learning rate was set to 1e-4 and weight decay was set to 1e-4.

In order to evaluate the performance of our model, we divided the Davis and GalaxyDB datasets to obtain the training, validation and test sets, respectively. And we divided the dataset according to the target proteins so that the target proteins in validation and test set were not seen in the training set. For the Davis dataset, the targets have the same sequence in train, validation and test sets are filtered out which results in 361 targets. Table 1 summarizes the split dataset in detail.

**Table 1.** Detailed information for the split dataset.

| Type | Proteins | Pos | Neg | Pairs |
| --- | --- | --- | --- | --- |
| Davis |  |  |  |  |
| Train | 231 | - | - | 15708 |
| Valid | 57 | - | - | 3876 |
| Test | 73 | - | - | 4964 |
| GalaxyDB |  |  |  |  |
| Train | 298 | 305702 | 1295867 | 1601569 |
| Valid | 38 | 43825 | 197666 | 241491 |
| Test | 36 | 31494 | 140505 | 171999 |

For the regression task on the Davis dataset, the performance of models was evaluated using Root Mean Square Error (RMSE), Mean Square Error (MSE), Pearson and Spearman. The main metric we used to evaluate the prediction performance in the classification task is the area under the receiver operation characteristic curve (ROC-AUC) metrics, which can reflect the ability of the model to correctly discriminate the active compounds and inactive compounds. And the AUC metric is also the condition for early stopping during model training. Additionally, we also measured the accuracy, recall, precision and F1 score metrics for evaluating the performance of the model prediction. It is worth noting that we determined the threshold for the above four metrics in the test set by finding the best threshold in the validation set. We set the search threshold in the range of 0.0 to 0.9, and searched with 0.001 steps to find the best threshold in the validation dataset according to the F1 score.

We compare our model with the following baselines:

1. SGDRegressor is a linear model fitted by minimizing a regularized empirical loss with Stochastic Gradient Descent (SGD). We used it for Davis dataset in the regression task. We experimented on the concatenated molecule and protein features. Here the molecule feature was Morgan Fingerprint calculated by RDKit [34], and the protein feature was the average tape embedding which suggests that taking the mean values at the amino acid level for original tape embedding.
2. L2-logistic regression (LR) applied a logistic regression model on the Morgan Fingerprint and tape embedding concatenated feature vectors, we used it for our GalaxyDB dataset in the classification task.
3. TransformerCPI [8] modified the transformer architecture with a self-attention mechanism to address sequence-based DPI classification task, we followed the default parameter settings in TransformerCPI and the same training and evaluating strategies as X-DPI.
4. GraphDTA [10] represented molecules as graphs and used graph neural networks to predict drug-target affinity. Here we compared our model with the GIN [37] in GraphDTA with default parameters. Besides, in order to fit the binary classification task on GalaxyDB dataset, we added a sigmoid function for the last layer in the GraphDTA network.
5. MolTrans [9] is an end-to-end biological-inspired deep learning-based framework that models the DPI process. We followed the same hyper-parameter setting described in the paper and compare our model with the MolTrans on our dataset.

## 3 Results and Discussion

### 3.1 Performance on Davis and GalaxyDB

In order to validate the effectiveness of our model, we first tested our model on a well-defined small dataset, Davis. As shown in Table 2, our model obtained the best MSE with 0.4738, which is 15.95% lower than the TransformerCPI model specifically and 5.30% lower than GraphDTA, the best performing baseline model. Interestingly, we finded that the performance of the complex deep learning models on the Davis dataset is not significantly better than other traditional machine learning models, and the strong learning ability of the deep learning model could not be well reflected on the Davis dataset. We believe this is due to the relatively small amount of data in the Davis dataset and the fact that it contains only proteins of the kinase family. We used this benchmark dataset to tune the model architecture.

**Table 2.** Performance comparison of our model and the baseline models on Davis dataset.

| Model | RMSE | MSE | Pearson | Spearman |
| --- | --- | --- | --- | --- |
| SGDRegressor | 0.7254 | 0.5262 | 0.5029 | 0.4483 |
| GraphDTA | 0.7073 | 0.5003 | 0.5519 | 0.4485 |
| MolTrans | 0.7233 | 0.5232 | 0.5332 | 0.4710 |
| TransformerCPI | 0.7508 | 0.5637 | 0.4414 | 0.3706 |
| X-DPI | **0.6883** | **0.4738** | **0.5552** | **0.4927** |

After the validation on a small dataset, we further evaluated the classification performance of different methods on the industry-scale GalaxyDB. Table 3 presented the overall performance comparison of our model and the baseline models, it noted that the threshold for the Precision, Recall and F1 score metrics in the test dataset was determined by finding the best threshold in the validation set as described in the previous section. And the final threshold corresponds to LR, GraphDTA, MolTrans, TransformerCPI and X-DPI are 0.191, 0.440, 0.412 and 0.449 respectively. Due to the best threshold for each model is different, the precision and recall would vary in different models.

**Table 3.**
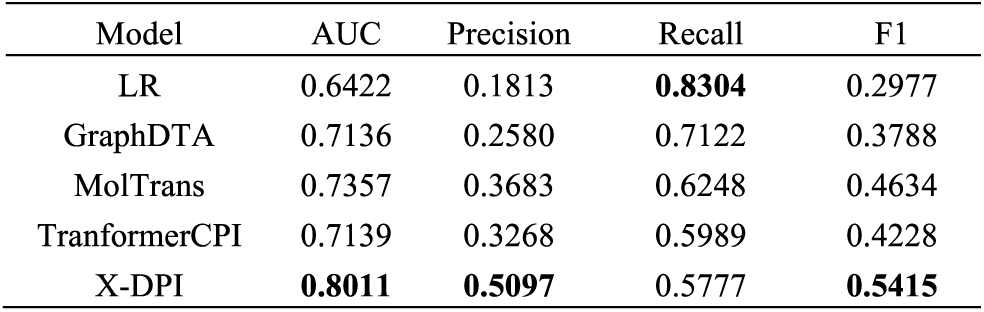
Performance comparison of our model and the baselines, where the precision, recall, and F1 score are calculated with the best threshold for each model.

| Model | AUC | Precision | Recall | F1 |
| --- | --- | --- | --- | --- |
| LR | 0.6422 | 0.1813 | **0.8304** | 0.2977 |
| GraphDTA | 0.7136 | 0.2580 | 0.7122 | 0.3788 |
| MolTrans | 0.7357 | 0.3683 | 0.6248 | 0.4634 |
| TransformerCPI | 0.7139 | 0.3268 | 0.5989 | 0.4228 |
| X-DPI | **0.8011** | **0.5097** | 0.5777 | **0.5415** |

In terms of overall performance, the proposed method achieved the best performance on AUC and competitive performance on other metrics compared to the baseline methods. It is noted that the best AUC is 0.8011, which is 8.89% higher than the best baseline MolTrans (0.7357) and 12.21% higher than the TransformerCPI (0.7139). The results showed that our proposed method has better prediction performance than other DPI prediction models. Figure 2(a) presented the receiver operating characteristic (ROC) curves comparison of all the methods in GalaxyDB test set. Figure 2(b) presented the precision-recall (PR) curves of all baseline models. We could seen that our model obtained consistent results and achieved superior performance over the other baseline models. This followed our expectations as the graph representation for proteins provides richer structure information than sequence representation, our model could provide more abundant information for molecules and proteins by combing the structure and sequence information from pre-trained embedding features, which could effectively improve the performance of DPI prediction. In order to validate the function of each component in our model, we ran several ablation experiments to analyze our model. Results were shown in Table 4 and we would discuss them in detail next.

**Table 4.**
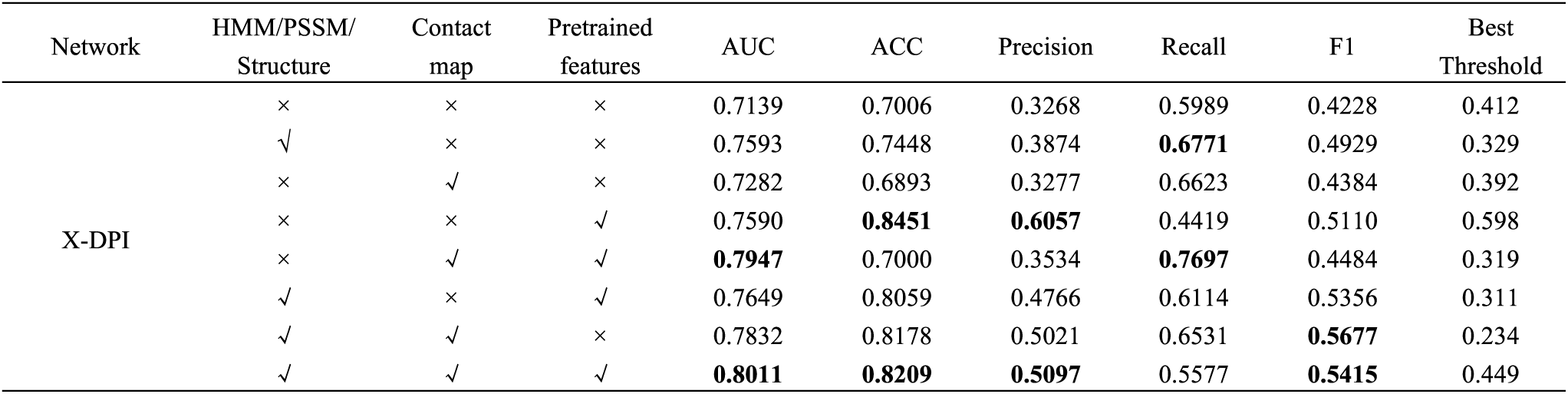
Results of ablation experiments.

| Network | HMM/PSSM/<br>Structure | Contact<br>map | Pretrained<br>features | AUC | ACC | Precision | Recall | F1 | Best<br>Threshold |
| --- | --- | --- | --- | --- | --- | --- | --- | --- | --- |
| X-DPI | × | × | × | 0.7139 | 0.7006 | 0.3268 | 0.5989 | 0.4228 | 0.412 |
|  | ✓ | × | × | 0.7593 | 0.7448 | 0.3874 | **0.6771** | 0.4929 | 0.329 |
|  | × | ✓ | × | 0.7282 | 0.6893 | 0.3277 | 0.6623 | 0.4384 | 0.392 |
|  | × | × | ✓ | 0.7590 | **0.8451** | **0.6057** | 0.4419 | 0.5110 | 0.598 |
|  | × | ✓ | ✓ | **0.7947** | 0.7000 | 0.3534 | **0.7697** | 0.4484 | 0.319 |
|  | ✓ | × | ✓ | 0.7649 | 0.8059 | 0.4766 | 0.6114 | 0.5356 | 0.311 |
|  | ✓ | ✓ | × | 0.7832 | 0.8178 | 0.5021 | 0.6531 | **0.5677** | 0.234 |
|  | ✓ | ✓ | ✓ | **0.8011** | **0.8209** | **0.5097** | 0.5577 | **0.5415** | 0.449 |

**Figure 2.**
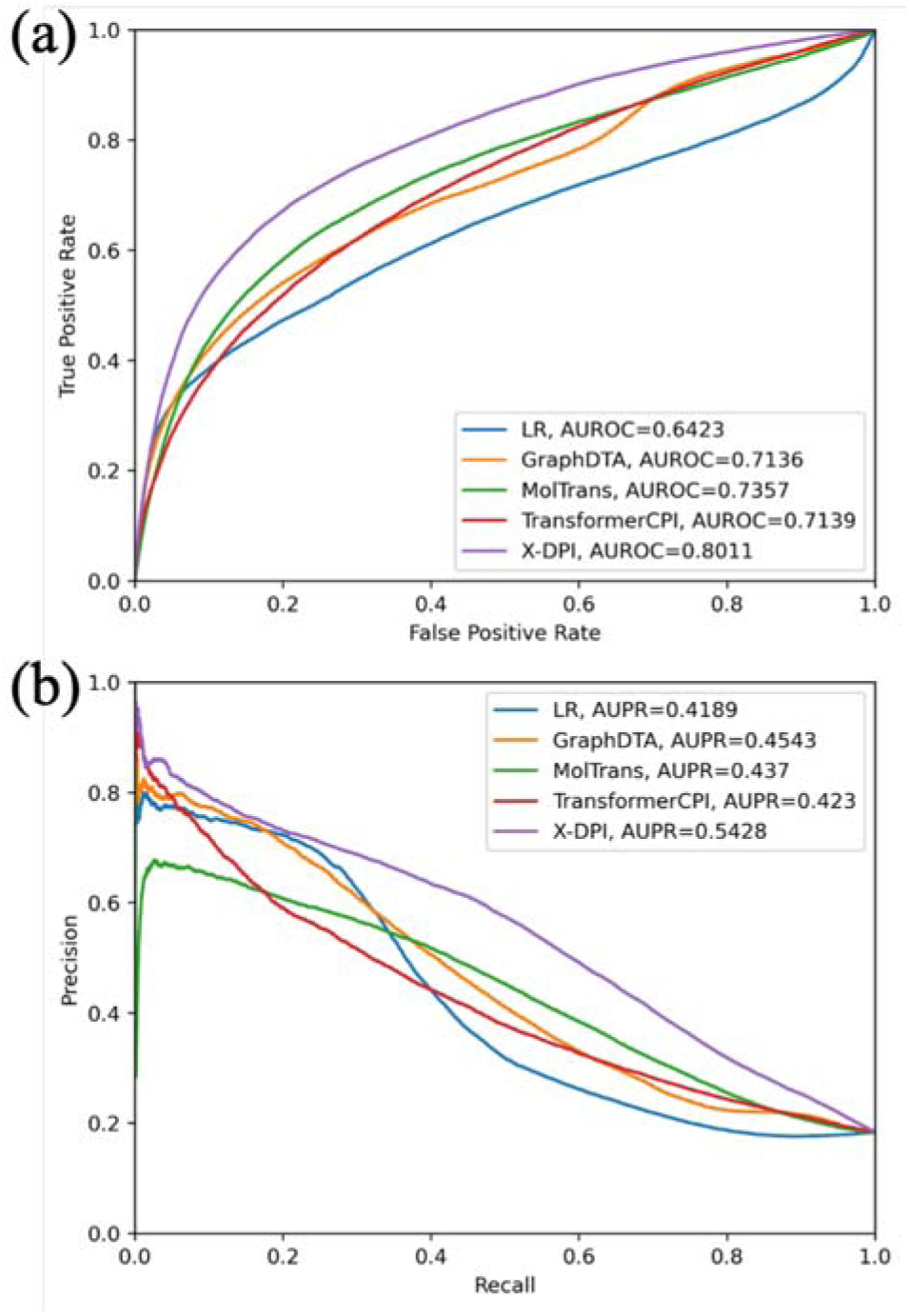
Performance of different methods on the GalaxyDB test set. (a) Receiver operating characteristice (ROC) curves of prediction results. (b) Precision-recall (PR) curves of prediction results.

### 3.2 Ablation experiments

Firstly, we evaluated the function of additional pre-trained embedding, which included the sequence information with tape embedding for proteins and substructure information for molecules. Instead of fusing the structure information from the protein graph and sequence information from tape embedding, we only used the structure information of proteins as the input features of the Transformer Decoder. We also used the structure information of molecules only in Transformer Decoder. As shown in Table 4, the removal of pre-trained embedding decreased the prediction performance of the model significantly. And this ablation experiment clearly showed the importance of pre-trained embedding in the model, which provides high-level protein sequence information and molecular substructure information for model learning.

Secondly, we evaluated the importance of structure information for protein in our model. In our model, the structure information of protein mainly came from the graph representation for protein, which included the amino acid node features HMM/PSSM/Structural features and the contact map of protein. In order to comprehensively evaluate the influence of protein structure information on model performance, we conducted ablation experiments on protein node features and contact maps, respectively. For the ablation experiments of the contact map, we avoided utilizing the protein structure information from the contact map and only use CNN to extract and fuse the node features in protein. The results represented in Table 4 suggests that the contact map processed by GCN could provide efficient and rich structure information for protein, which results in a better performance in the DPI prediction task. However, due to the structure of proteins is extremely complex that the predicted contact map can’t accurately reflect the structure of proteins, we still need to introduce additional information such as evolutionary and predicted structural features to compensate for the information loss of the predicted contact map. And the ablation experiments have also shown that our model could significantly improve the performance by combing the contact map and other additional protein information. To validate the function of protein node features in our model for DPI prediction, we used word2vec embedding in TransformerCPI to replace the protein node features as HMM, PSSM and structural features. We observed that when we used the word2vec embedding as the protein node feature, the performance has slightly decreased. This suggests that the HMM, PSSM and structural features used in our model can provide better protein information at the amino acid level compared to word2vec, which is beneficial to the accurate prediction of the DPI prediction task.

### 3.3 Performance on the prospective validation

In order to verify the generalization ability of our model, we used the model trained on the GalaxyDB training set to evaluate the Davis dataset and external test set from AstraZeneca.

For the Davis dataset in prospectives validation, the experimental results were shown in Figure 3. In general, our model consistently performed well in the test set. When the test proteins were observed in the training set, X-DPI achieved an AUC of 0.8813, which is 5.28% higher than the best baseline GraphDTA (0.8371) and 8.23% higher than the TransformerCPI (0.8143). For the unseen proteins, our model also achieved the best AUC with 0.7073, which is 4.77% higher than GraphDTA (0.6751) and 11.21% higher than TransformerCPI (0.6360).

**Figure 3.**
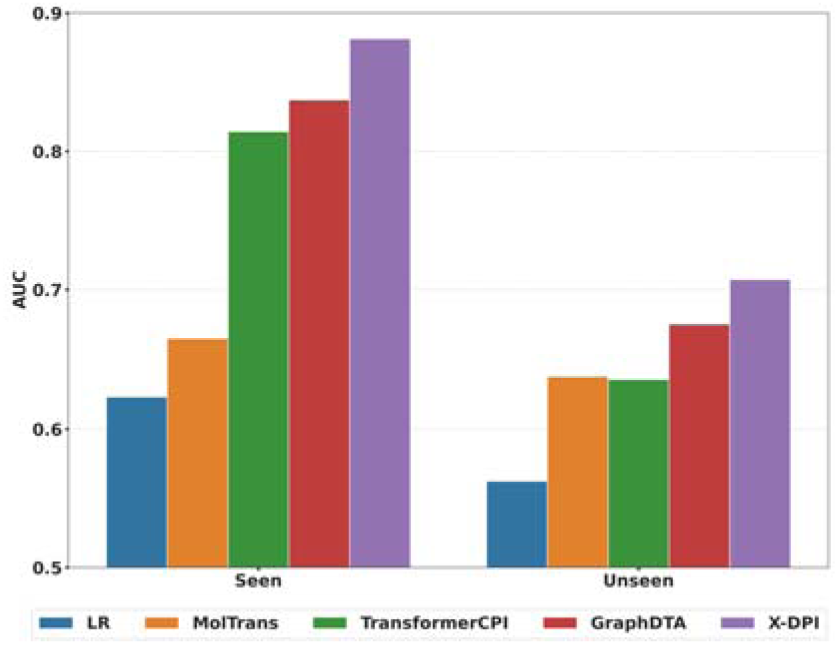
Performance comparisons of X-DPI and baselines on seen and unseen protein targets in the Davis dataset.

For the external test dataset in prospective validation, the data distribution and classification performance of our model were given in Figure 4. As shown in Figure 4, for the seen proteins, the average AUC on a total of 30 targets is 0.7724, and 70% of the seen targets reached AUC≥ 0.7. For the unseen proteins, the average AUC on a total of 10 targets is 0.7264 and 50% of the seen targets reached AUC≥ 0.7.

**Figure 4.**
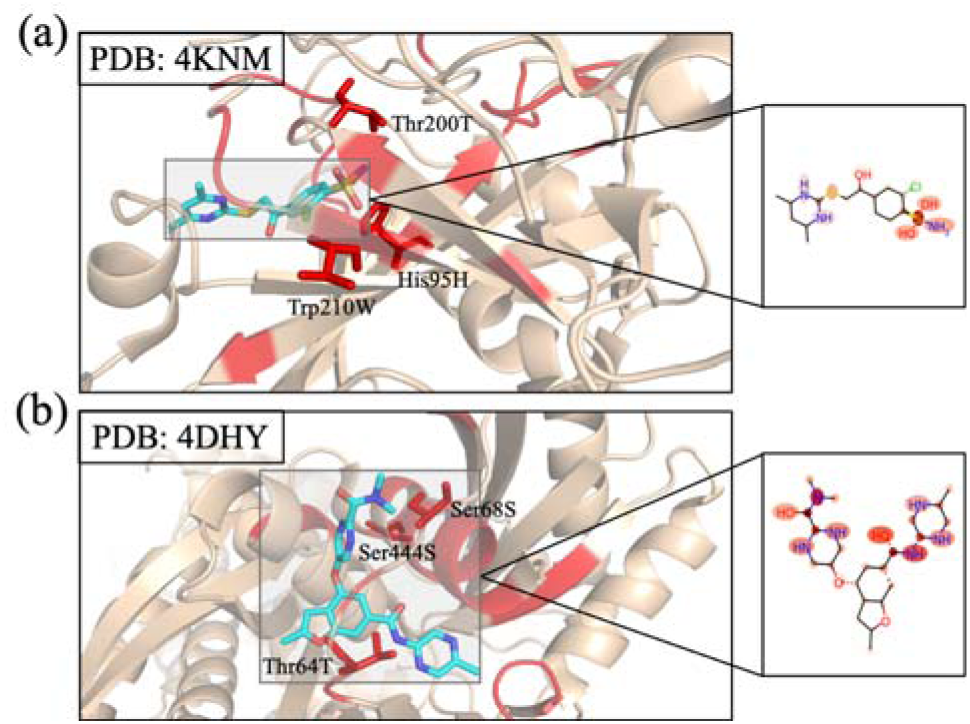
The information and evaluation results about the external test dataset. (a) The data points distribution of individual target proteins. (b) The AUC performance of our model on an external test dataset. The box plot represents the AUC distribution of individual target protein performance for seen and unseen proteins.

In general, our model achieved reasonable performance on both seen and unseen proteins, indicating that the X-DPI trained on GalaxyDB generalizes well to independent virtual screening tasks. However, performance gaps between seen and unseen proteins were observed both on the Davis dataset and external test dataset. We argue that there might be two potential reasons for these performance gaps. The first one is that the chemical space for seen and unseen target proteins is different, which makes the knowledge of chemical spatial distribution for seen proteins learned by our model unable to be effectively applied to unknown chemical space for unseen proteins. The second is that the ability of our model to learn unseen protein representations is still somewhat deficient. The predicted contact maps and protein pre-trained embeddings have helped us to improve our predictions for unseen proteins, but there is still much room for improvement.

### 3.4 Study for Interpretability

Thanks to the Transformer Interaction Decoder architecture module, our model is able to analyze the interaction mechanism between the protein and the molecule. The positions focused on the self-attention mechanism can provide a reasonable explanation for the binding activity prediction, and also help the researchers to quickly locate the key interaction sites between protein and molecule when performing further activity analysis.

To exemplify this, we selected two complexes from RCSB Protein Data Bank (PDB) [38] as the representatives, where the proteins are presented in the test of GalaxyDB. In particular, we colored the top-weighted residues of the example proteins and atoms of the ligand with red and compared them to the actual protein-ligand interaction sites retrieved from the PDB. We found that the highest-weighted amino acids and molecular atoms overlap substantially with the real interaction sites. For protein CA13 (UniProt ID: Q8N1Q1) in Figure 5(a), the attention bar highlights residues His95H, Thr200T and Trp210W, which highly overlap with the key pocket residues observed in the co-crystal complex (PDB: 4KNM). For protein GCK (UniProt ID: P35557) in Figure 5(b), the highlighted residues (Thr64T, Ser68S, Ser444S) and ligand functional groups in the importance maps show high similarity to observed interactions in the co-crystal complex (PDB: 4DHY). The results suggest that the model can be applied to analyze the interaction mechanism between molecules and target proteins and inspire researchers.

**Figure 5.**
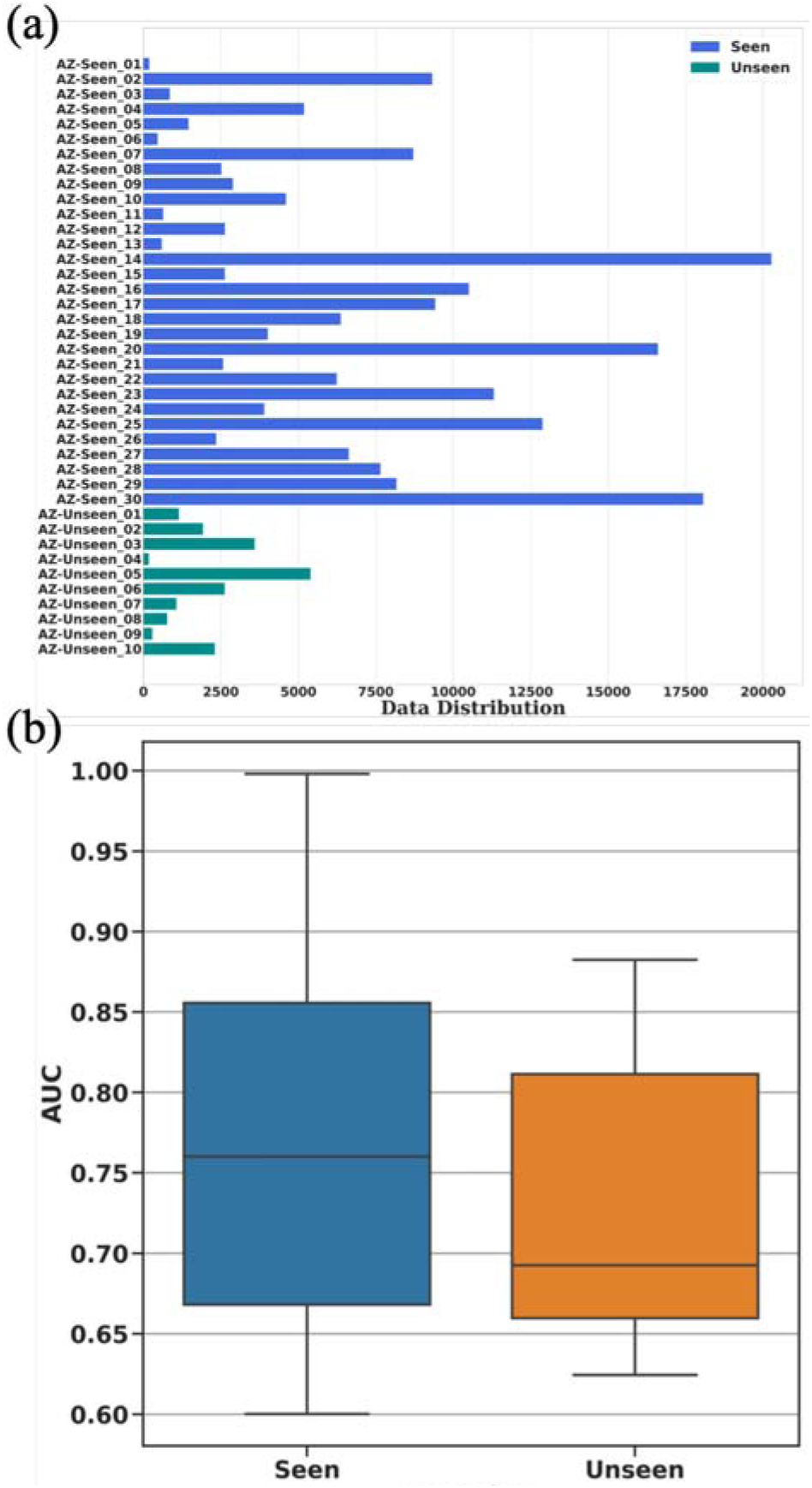
Attention weights visualization of pocket and ligand pairs. (a) Attention weight of interaction for 4KNM and E1E. (b) Attention weight of interaction for 4DHY and S41.

## 4 Conclusion

In this study, we selected the Davis and GalaxyDB dataset as the internal validation dataset for our model, meanwhile, we further verified the generalization ability of our model on the external test set collected from AstraZeneca. For the GalaxyDB dataset, we first analyzed the data distribution of the ExCAPE-ML dataset for the DPI prediction task, then screened the dataset to generate a high-quality benchmark dataset named GalaxyDB, which is a more balanced dataset for the distribution of positive and negative samples than ExCAPE-ML and has enough data to effectively train and evaluate the deep learning model. At the same time, we proposed a DPI prediction model named X-DPI with better performance than other baseline models. We compared our model with the previously reported models for the DPI prediction task, and the experimental results have shown that our model achieved the best MSE with 0.4631 on Davis and best AUC with 0.8011 on GalaxyDB in all baseline models. In order to enable the model to extract protein structure information, we tried to use graph structures to represent the proteins. In the protein graph, we used the predicted contact map of a protein to represent the adjacency matrix and treat each amino acid in the protein as a graph node. Meanwhile, we used evolutionary features like HMM, PSSM and structural features for proteins as the node features in the protein graph. Besides, we tried to use the pre-trained embedding in our model to increase the high-level similarity information of instances for molecules and proteins. We utilized the sequence features represented by tape embedding for protein sequences and substructure features represented by mol2vec for molecules. Finally, we obtained a better feature representation for proteins and molecules by combining the structure and high-level information in our model, which leads to a better performance for DPI prediction in the benchmark dataset. The ablation experiments have shown that the protein graph constructed by the contact map and amino acid node can provide richer and more accurate structure information for DPI prediction, and we could obtain better performance for DPI prediction when we combine high-level pre-trained information from the protein and molecules.

Overall, we believe that our study provides a new SOTA model for DPI prediction research. Additionally, the benchmark dataset that we constructed can be used for the community to develop and evaluate future DPI models. Combining the structure and pretrained information for both proteins and ligands seems provides advantages in making DPI prediction and could be a new area to explore in the future..

## Key Points

- We curated a high-quality benchmark dataset named GalaxyDB specifically designed for machine learning-based virtual screening. GalaxyDB was derived from ExcapeDB and consists of 372 common targets with 381,021 confirmed active and 1,634,038 confirmed inactive compounds. The large-scale dataset and an unbiased training procedure provide advantages for model building than using a small toy dataset.
- For informative protein representation, we represented the protein sequence as a protein graph by combining the predicted contact map, evolutionary and secondary structural features, meanwhile, we constructed a structure-aware graph neural network method from the protein sequence by combining protein graph and graph neural networks.
- We introduced self-supervised pre-trained embedding of drugs and proteins, respectively, in order to strengthen the protein/drug association signals. Our model leverages this high-level information in a unified framework and generates interpretable results with a transformer-based interaction mechanism.

## Biography

Penglei Wang is a master student at Institute of medical Robotics at the Shanghai JiaoTong University. His research interests lie in deep learning and drug discovery.

Shuangjia Zheng is a PhD student in Computer Science and Technology at the Sun Yat-Sen University. His research interests lie in deep learning, knowledge graph embedding, drug discovery and computational biology.

Yize Jiang is a senior algorithm engineer in Galixir, Beijing, China, he received his master degree from the Beijing Institute of Technology. His research interests are drug discovery, computational biology and machine learing.

Chengtao Li is the Chief Executive Officer of Galixir, Beijing, China, he received PhD degree from the Massachusetts Institute of Technology. His research interests are drug discovery and machine learning.

Junhong Liu is a senior algorithm engineer in Galixir, Beijing, China, he received his master’s degree from the Peking University. His research interests are drug discovery, computational biology, and machine learning.

Chang Wen is a research assistant in Guangzhou Regenerative Medicine and Health Guangdong Laboratory, Guangzhou, China. Her research interests are computational biology and data mining.

Atanas Patronov is a senior data scientist in MolecularAI, Discovery Sciences, BioPharmaceuticals R&D, AstraZeneca, Gothenburg, Sweden. His research interests are drug discovery and data mining.

Dahong Qian is a Professor at the Institute of Medical Robotics, Shanghai Jiao Tong University, Shanghai, China. His research interests are clinical-driven AI-based computer-aided diagnostics and therapeuritcs, microdevices and embedded AI algorithms for medical robotics, and wearable and implantable devices.

Hongming Chen is a Principal Investigator in Guangzhou Regenerative Medicine and Health Guangdong Laboratory. His research interests are computational chemistry and computational biology.

Yuedong Yang is a Professor in the School of Data and Computer Science and National Super Computer Center at Guangzhou, Sun Yat-sen University, China. Currently, his research group emphasizes developing HPC and AI algorithms for multi-omics data integration and intelligent drug design. He is also responsible for constructing the HPC platform for biomedical applications based on the Tianhe-2 supercomputer.

## Funding

This study has been supported by the National Key R&D Program of China (2020YFB020003), National Natural Science Foundation of China (61772566), Guangdong Key Field R&D Plan (2019B020228001 and 2018B010109006), Introducing Innovative and Entrepreneurial Teams (2016ZT06D211), Guangzhou S&T Research Plan (202007030010).

## Conflict of Interest

We declare competing interests. This work is done when P.W works as intern at Galixir. S.Z., Y.J, C.L. and J.L were employees of Galixir, and A.P was employee of AstraZeneca.

## Acknowledgments

We thank the Galixir team for its support and discussion, and with special thanks to Jixian Zhang, Zixuan Liu and Da Wei for the experimental design discussion and technical support.

## Availability and implementation

The datasets and code will be available soon.

